# Loss of Stk11 in basolateral amygdalar projection neurons impairs taste aversion learning by altering the temporal pattern of taste response plasticity in gustatory cortex

**DOI:** 10.1101/2021.05.31.446460

**Authors:** Roshan D. Nanu, Thomas J. Murdy, David Levitan, Seneca Scott, Sacha B. Nelson, Donald B. Katz

## Abstract

Gustatory cortex (GC) responds to tastes on the tongue with dynamic ensemble activity that represents first the presence, then the identity, and finally the hedonic value (palatability) of tastes. This final state of the taste response is uniquely altered by conditioned taste aversion (CTA) – a powerful one-trial learning paradigm in which a taste becomes aversive after association with gastric malaise – a process requiring coordination between GC and basolateral amygdala (BLA). Previously, we found that one key requirement for learning in this circuit is expression of the serine/threonine kinase 11 (Stk11) gene (a tumor suppression gene which has only recently been associated with learning; Levitan et al., 2020). When Stk11 is knocked out in BLA projection neurons (BLApn), CTA learning fails to occur. Here we have examined how learning-related response plasticity in GC taste responses is impacted by the knockout of Stk11 in BLApn. Contrary to the commonly held assumption that a lack of learning means a lack of such plasticity, but consistent with the fact that Stk11KO has been shown to increase the excitability of BLApn, our data reveal that the knockout of Stk11 in BLApn does not eliminate plasticity; rather, it shifts the impact of CTA training on GC taste responses to an earlier, learning-inappropriate epoch. Even naïve taste representations are altered—specifically, the pattern of similarities and differences among the different taste responses are rendered abnormal by Stk11 KO, and these relationships fail to change with training. Finally, the latency of behavior-related dynamic ensemble features of the GC taste response, which is also abnormal in naïve KO mice, is rendered disorganized by CTA. Together, these results suggest that Stk11 plays a role in governing the coordination of GC activity by BLA, and demonstrate that alterations in the function of BLApn caused by Stk11KO inhibit learning not by inhibiting plasticity but by changing its temporal properties.

## Introduction

Following placement of a taste on the tongue, gustatory cortical (GC) ensemble responses exhibit a rich temporal structure, evolving through a sequence of distinct encoding states that signal first the presence of a taste, then the identity of the taste, and finally the palatability or hedonic value of the taste (Katz et al., 2001; Jones et al., 2007; Piette et al., 2012; Sadacca et al., 2012). On individual taste presentations, the onset of the palatability state is a sudden and coherent ensemble transition, the timing of which reliably predicts and drives behavior (Moran & Katz, 2014; Sadacca et al., 2016; Mukherjee et al., 2019).

When presentation of a particular taste is paired with induced gastric malaise such that an animal learns a conditioned taste aversion (CTA), response plasticity is temporally-specific: the coding content (as measured in firing rates) of the palatability phase of that taste’s response changes to reflect depreciation of the tastant’s value, while pre-training activity in prior response phases is largely preserved (Grossman et al., 2008; Moran & Katz, 2014; Lavi et al., 2018).

This learning-relevant palatability-related information appears to be in part provided to GC by the basolateral amygdala (BLA) (Piette et al., 2012; Lin et al., 2021). BLA is necessary for proper CTA learning (Gallo et al., 1992; Yamamoto et al., 1995; Grossman et al., 2008; Garcia-Delatorre et al., 2014), as are the axons of BLA→GC projection neurons (BLApn) themselves (Yamamoto et al., 1984; Levitan et al., 2020). Of particular interest in this regard are recent findings demonstrating that the BLApn-specific knockout of the serine/threonine kinase 11 (Stk11) gene, commonly studied as a tumor suppressor and regulator of cell growth in tissue throughout the body (Shaw, 2009; Nakada et al., 2010; Beirowski, 2019), prevents the acquisition of CTA (Levitan et al., 2020).

At the cellular level, the effect of losing Stk11 is an increase in the basal excitability of BLA→GC projection neurons (Levitan et al., 2020), a fact that has intriguing implications for the impact of the knockout on taste-related GC neural activity over CTA training. Most generally, these results suggest the possibility that the Stk11 KO may cause an “over-drive” of GC by BLA input; far from inhibiting plasticity, the increase in excitability might be expected to enhance GC response plasticity while rendering it temporally inappropriate for learned behavior.

Here, we have set out to test this unusual hypothesis, investigating how the knockout of Stk11 in BLApn disrupts learning-related changes in GC taste responses. We recorded extracellular single neuron responses in GC of awake, freely behaving mice as these mice underwent CTA training, and observed how taste responses in GC changed—both those of individual neurons and of ensembles of neurons. Our results reveal subtle alterations of taste processing in naïve Stk11 KO mouse, and show that, in these mice, CTA training-related changes are not nonexistent but rather learning-inappropriate.

## Results

We performed a selective, inducible knock out of Stk11 in BLA projection neurons (Stk11 KO, N=6 mice) using viral transfection to express Cre in Stk11^fl/fl^ mice. One week later, we implanted bundles of electrodes bilaterally in GC and inserted an intraoral canula (IOC) under the masseter muscle so that tastes could be delivered directly onto the tongue. Ten days after virus injection and electrode/IOC implantation, we recorded bilaterally from ensembles of GC single neurons throughout a combined tasting/CTA training regime: mice were presented with a battery of tastes of known hedonic values; one day later, the mice were subjected to a 24hr CTA protocol designed to train an aversion to saccharin, a taste that is palatable to the naïve mouse, and then given another, post-learning taste battery (Figure 1B). We also subjected a control cohort (Control, N=10 Stk11 ^fl/fl^ mice) that had undergone injection of a viral vector coding only for GFP (Figure 1A) to the same experimental regime. Given that CTA learning occurs across <24hrs (with only one taste-illness pairing), we were able in some cases to monitor the responses of single neurons in GC from before until after learning.

**Figure 1.**
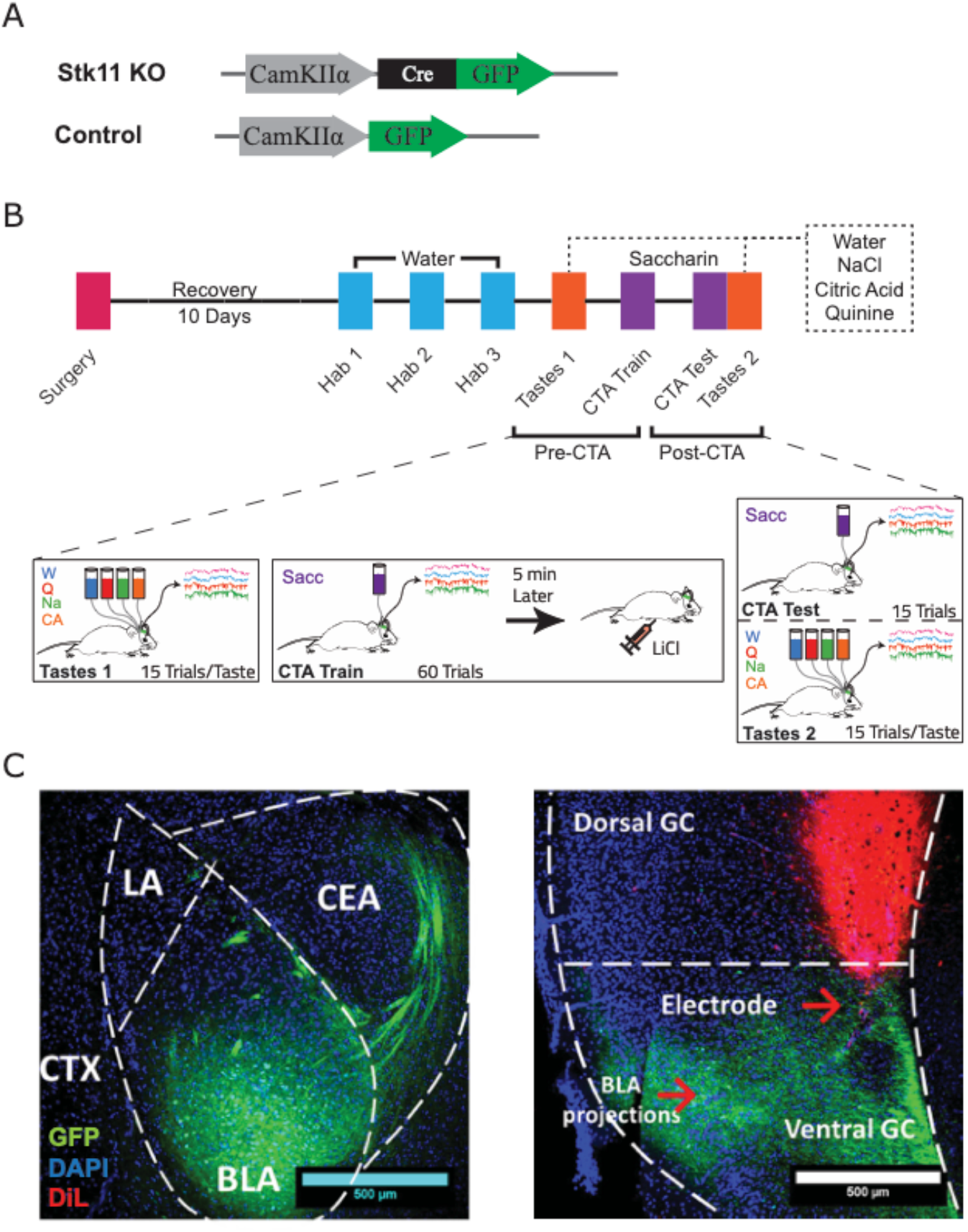
Overview of experimental setup. (A) Genetic constructs Cre-GFP (Stk11 KO) and GFP (Control) under the control of a CamKII*α* promoter were encapsulated in AAV2/5 viral vectors. (B) Overview of the experimental timeline starting from the virus injection and electrode/IOC implantation surgery through to recording days – Tastes 1, CTA Train, CTA Test and Taste 2. Except for the 10-day recovery, the spacing between blocks indicates the passage of 1 day. (C) Representative histological slices showing viral transfection of BLA (left; −1.2mm AP) and co-localization of electrodes (red, DiL stain) and BLA axon terminals (green, GFP) in ventral GC (right; +1mm AP).

We evaluated learning by comparing nightly water bottle consumption before CTA training to saccharin consumption after CTA training (10 min access each time). Given the complexity of the recording protocol (see Methods), some extinction was expected to occur before the test of learning. Furthermore, relatively mild learning is an inevitable result of embedding the training in a context of multiple recording sessions involving presentation of multiple tastes (see Discussion). In recognition of these factors, we set a relatively low threshold: animals whose saccharin consumption was <80% of their mean nightly water consumption were deemed to have learned and retained the aversion; this criterion was reached by 5 of the 10 control mice, but by none of the Stk11 KO mice that drank water normally (see Methods). These results are consistent with those shown previously (Levitan et al., 2020)—Stk11 knockout in BLApn inhibits normal CTA learning.

A total of 460 well-isolated single neurons (313 from Stk11 KO mice; 147 from control mice that learned CTA) were subjected to initial analysis (13 ± 2.1 cells per session). The basic properties of taste responses were largely similar for neurons from the two groups of mice, but Figure 2 (top-left) shows that the average firing rates of putative pyramidal neurons, computed over the 1s before stimulus presentation (baseline), were higher in Stk11 KO mice than in control mice (95% CI of difference: [0.40, 1.0] Hz). A two-way ANOVA comparing group and training (pre-vs post-CTA) effects on pyramidal cell firing rates revealed a significant group effect (F(1,384)=5.1, p=0.024) on baseline firing rates (no other effects were significant). No between-group differences were observed between interneuron firing rates, and whole-response firing rates across the 1.5s after taste delivery (response, with no examination of “temporal coding”) were similar between groups for both cell types.

**Figure 2.**
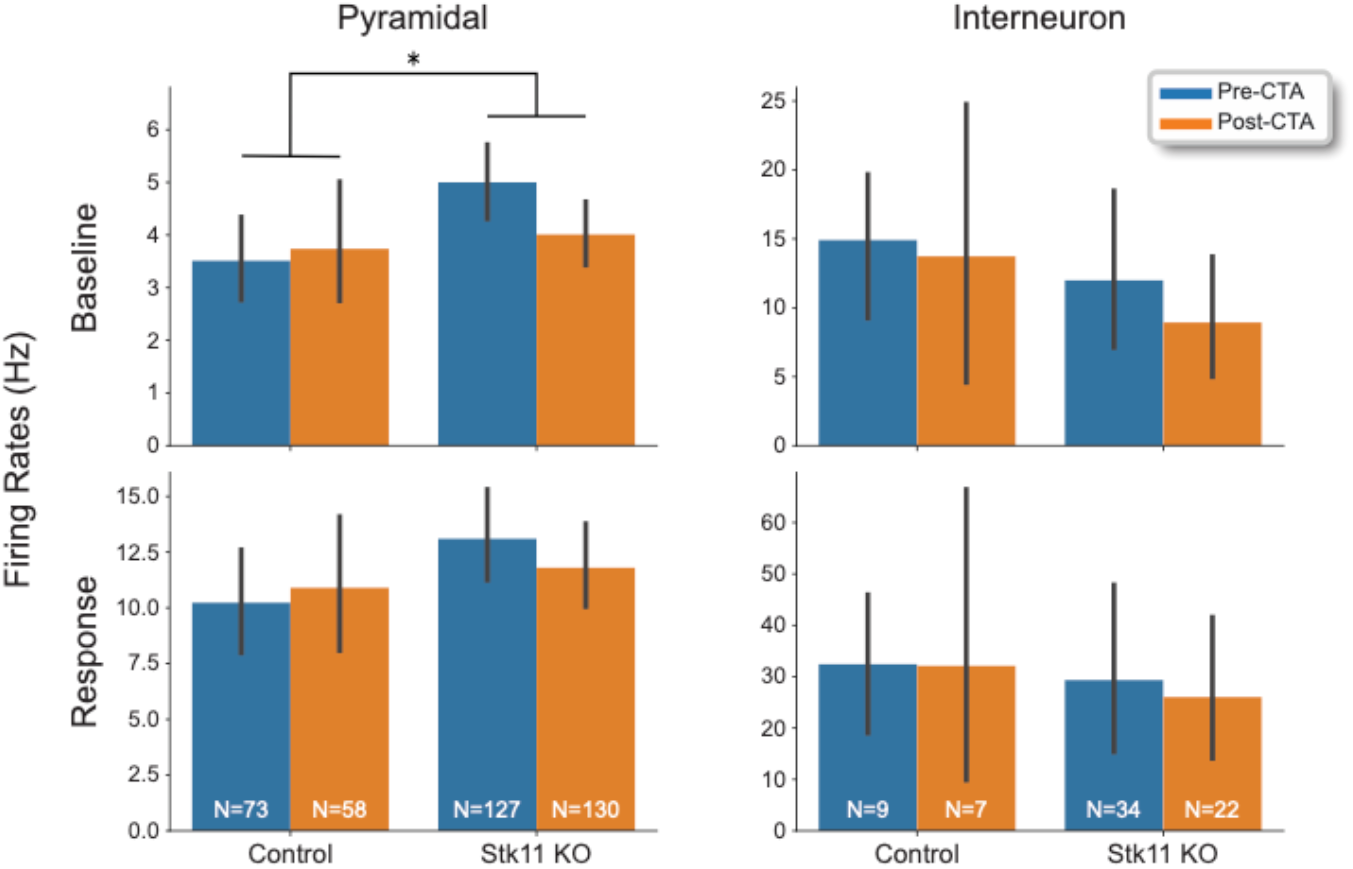
Pyramidal neurons in Stk11 KO mice exhibit elevated baseline firing rates. Baseline (top; 1s pre-stimulus) and response (bottom; 1.5s post-stimulus) firing rates for pyramidal cells (left) and interneurons (right) in both groups over CTA training. Error bars show the 95% confidence intervals (CI) for the means. Pyramidal cell basal firing rates were slightly (95% CI of difference: [0.396, 1.03] Hz), but significantly, elevated in Stk11 KO mice. *p<0.05.

### The dynamics of learning-related response plasticity is abnormal in the BLA-specific Stk11 KO

In GC responses, palatability-correlated firing emerges across the 1.5s after stimulus delivery (Levitan et al., 2019; Levitan et al., 2020). The same pattern was observed here (Figure 3A) for both groups of mice. A two-way ANOVA for group and training effects performed on these palatability correlation data revealed significant effects of both the KO (F(1,1940)=63, p=3.0e-15) and training (F(1,1940)=6.4, p=0.012; Figure 3B)—Stk11 KO mice demonstrated a consistently higher degree of palatability-related correlation than Controls (95% CI of difference: [0.029, 0.044]), and both groups showed a mild elevation in palatability correlation after training.

**Figure 3.**
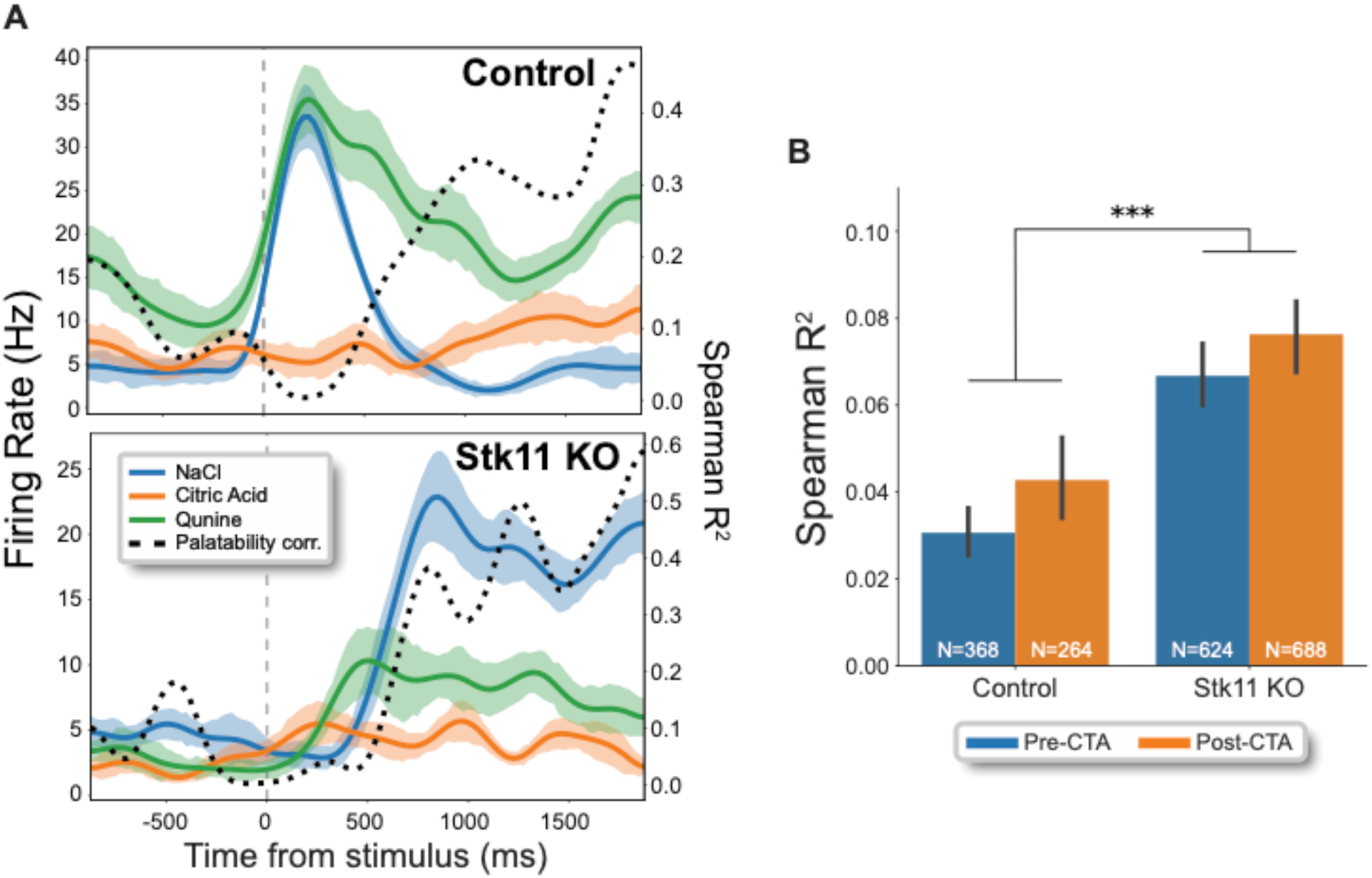
Enhanced correlation to taste palatability in single neuron taste responses of Stk11 KO mice. (A) Peri-stimulus time histograms of representative single neurons from control (top) and Stk11 KO (bottom) mice in response to tastes from the Tastes 1 (pre-CTA) session: NaCl (blue), Citric Acid (orange) and Quinine (green); water was excluded since it’s palatability could not be properly ranked relative to the other stimuli. The shaded regions show the SEM, and the dashed black line indicates the Spearman R^2^ correlation coefficient (right axis) for the unit firing rates to the rank-order palatabilites of the tastants; note that for both, the palatability-relatedness of the firing increased across the 2^nd^ half-second of the response. (B) Average correlation values across 1.5 sec post-stimulus for all cells. Error bars represent the 95% CI of the means. The palatability-relatedness of GC firing was enhanced in Stk11 KO animals compared to controls (95% CI of difference: [0.0286, 0.0435]); ***p<0.001.

In our previous paper (Levitan et al. 2020), we failed to observe significant differences in the correlation of GC firing to palatability between Stk11 KO and Control. This small discrepancy between these data and our prior results is likely due to our increased sample of Stk11 KO neurons and the fact that our current taste battery lacked sucrose, the reliable high palatability of which may have overwhelmed subtle between-group differences (see Discussion). Regardless, the slight, general enhancement in GC palatability-related firing makes sense: since a portion of palatability information in GC appears to come from BLA projections (Piette et al., 2012; Lin et al., 2021), a larger than normal contribution of BLA to GC taste responses might be expected resultant from the increased BLA excitability in the Stk11 knockout (Levitan et al., 2020).

The fact that CTA training changed GC responses in Stk11 KO mice (mice that failed to learn CTAs) was more surprising, as failure to learn is typically assumed to imply a lack of training-related response plasticity. CTA learning, however, is supported not simply by response changes, but by changes that are specific in timing and nature—changes in late-epoch firing that leave the conditioned taste responses more similar to those caused by innately aversive tastes (Grossman et al., 2008; Moran & Katz, 2014; Lavi et al., 2018; Mukherjee et al., 2019; Arieli et al., 2020). We therefore hypothesized that GC response changes wrought in Stk11 KO mice by CTA training might differ from those observed in control mice in these temporal specifics. To test this hypothesis, we focused on evaluating stimulus-evoked response dynamics in neurons held across the entire CTA training regimen. Firing rates relative to baseline were compared for the 1.5s post-stimulus in 250ms non-overlapping bins (and statistically analyzed using one-way ANOVA with Bonferroni correction).

As has been shown previously for rats (Grossman et al., 2008; Moran & Katz, 2014), significant learning-related changes in GC neurons recorded from control animals were largely restricted to the late epoch of the response (>750ms; Figure 4a, left). This pattern is consistent with the generally agreed-upon suggestion that CTA changes the palatability of a taste, leaving taste identity intact (Bures et al., 1998; Fonseca et al., 2020), and with the fact that it is the transition into this late state that is implicated in the driving of palatability-related behavior (Li et al., 2016; Sadacca et al., 2016; Mukherjee et al., 2019).

**Figure 4.**
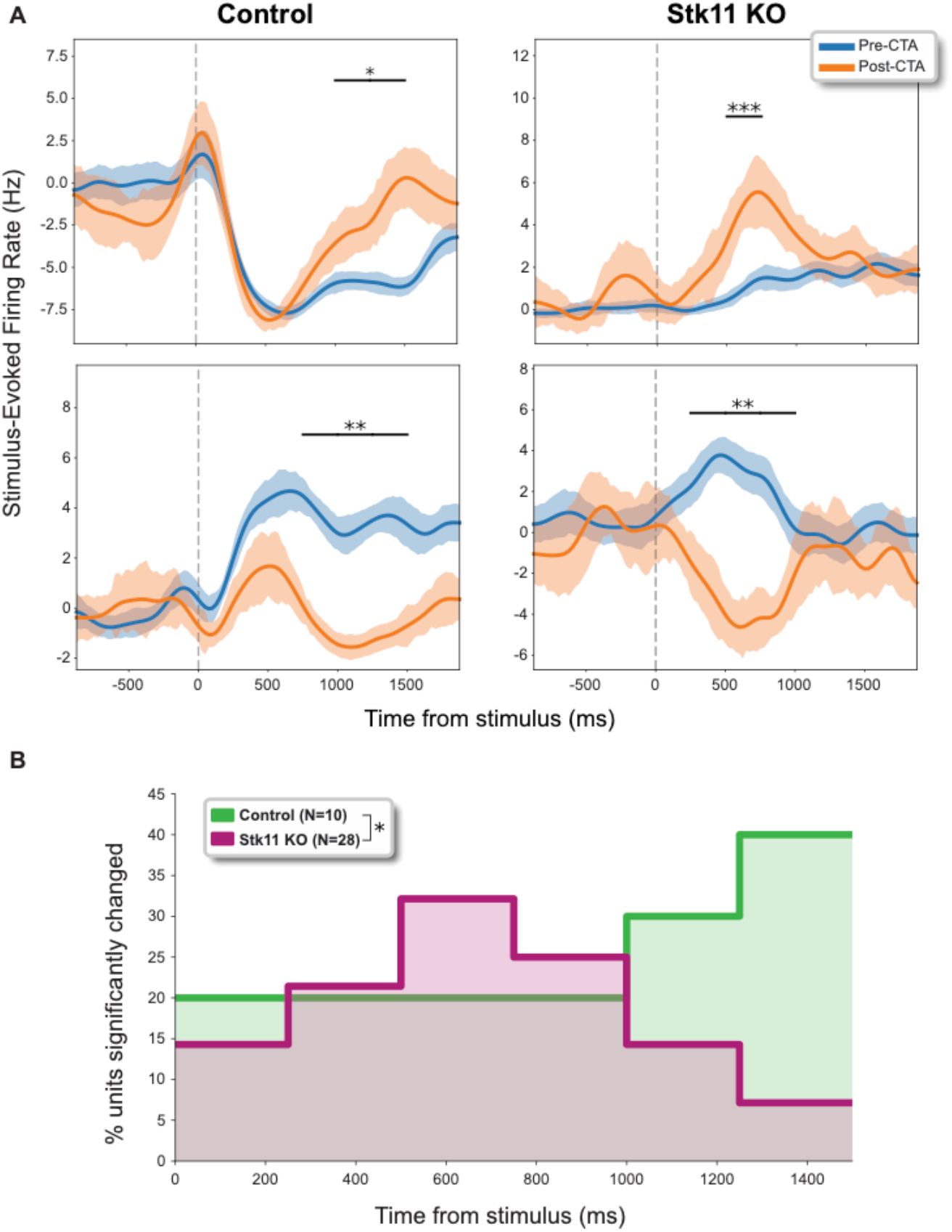
CTA training causes earlier changes to single-neuron saccharin responses in Stk11 KO mice than controls. (A) Saccharin-induced firing rates (relative to mean baseline) before and after CTA for representative single neurons from control (left) and Stk11 KO (right) mice. Asterisks with horizontal lines identify significant differences — the results of one-way ANOVAs conducted on each time bin and Bonferroni corrected. (B) Percent of single neurons held across training and testing sessions for which training significantly changed firing rates, for each time bin. For Stk11 KO mice (red), changes were found in earlier portions of the responses than was true for control animals (green); *p<0.05, **p<0.01, ***p<0.001.

These findings contrast with the results for Stk11 KO mice. Training-related changes were in fact present in these mice, but the response plasticity was abnormally centered on early parts of the response, with firing rates returning to pre-training levels by the late phase (Figure 4A, right). Figure 4B shows the examples in Figure 4A to be representative: training-related response plasticity in the control group responses tended to appear late in responses, but clustered at earlier response times in Stk11 KO mice; these distributions differed significantly (χ^2^(1)=13, p=0.026, V=0.580), confirming that training-related response plasticity in Stk11 KO mice, while robust, was abnormal.

It was important to consider the possibility that this seemingly abnormal response plasticity observed in Stk11 KO mice was not simply a “normal” result of the fact that these mice failed to learn. Prior work from our lab suggests that this is not the case, in that sham-CTA, in which no aversive element is introduced, results in almost no (<5% of cells) response plasticity (Grossman et al., 2008). Additionally, examination of GC responses in the subset of control mice for which learning failed to reach significance (data not shown) revealed a distribution that looked, if anything, more like those of control mice that learned (χ^2^(1)=2.1, p=0.83, V=0.36) than they did Stk11 KO mice (χ^2^(1)=14, p=0.018, V=0.63)—a fact that likely reflects learning rendered subthreshold by extinction and the inclusion of a taste battery (see Discussion). Thus, we conclude that the abnormal training-related taste response plasticity observed in our Stk11 KO mice is not the result of a normal failure or absence of learning, but truly reflects a change in training-related plasticity wrought by the Stk11 KO in BLApn.

### Impaired ensemble representations of saccharin with Stk11 KO

The above analyses demonstrate that CTA training changes the content of GC taste responses in both control and Stk11 KO mice, but that the nature of those changes (defined by known differences in coding in different temporal epochs of the taste responses, see Fontanini & Katz, 2006; Jones et al., 2007; Miller & Katz, 2010; Maffei et al., 2012; Moran & Katz, 2014; Sadacca et al., 2016) is group-dependent; the plasticity is learning-appropriate in control mice only. These analyses beg the question of the implications of these changes for the information content of the responses.

To investigate this aspect of response plasticity, we examined the information available in the subset of single neurons that were held across either both pre-CTA or both post-CTA recording sessions so that saccharin responses could be related to those of other tastants. We embedded the population firing rates from the early (<750ms) and late (750-1500ms) epochs of the response into separate 2-D spaces using the manifold data reduction technique multi-dimensional scaling (MDS). By taking the Euclidean distances between saccharin responses and the mean quinine and NaCl responses (distances D_Q_ and D_N_, respectively), and normalizing to the distance between the mean quinine and NaCl responses (D_QN_), we created a dimensionless distance metric (*D*_*S*_=[*D*_*Q*_ − *D*_*N*_]/*D*_*QN*_) that allowed us to accurately compare how similar the ensemble saccharin representations (even those with different numbers of neurons) were to each. Values greater than 0 signified that the saccharin representation was more similar to NaCl (the palatable tastant) than to quinine. Note that this analysis does not necessarily reflect the amount of palatability information embedded in the ensemble response, but instead shows the organization of the N-dimensional saccharin representation in relation to those of tastants whose valence remains fixed over the CTA paradigm (see Methods).

The results of this analysis are presented in Figure 5. Consistent with what is known of CTA learning (Moran & Katz, 2014), late epoch saccharin representations in control mice shifted toward those of quinine as a result of CTA training (see Table 1). Saccharin responses in the Stk11 KO group, meanwhile, showed no such change. Instead, these responses were significantly closer to quinine than those of controls before training, which then failed to change that coding significantly; if anything, the “early epoch” responses (which were the site of most of the training-related changes, see Figure 4b) became more “NaCl-like” with training – an opposite change than that expected to occur with CTA learning.

**Table 1.**
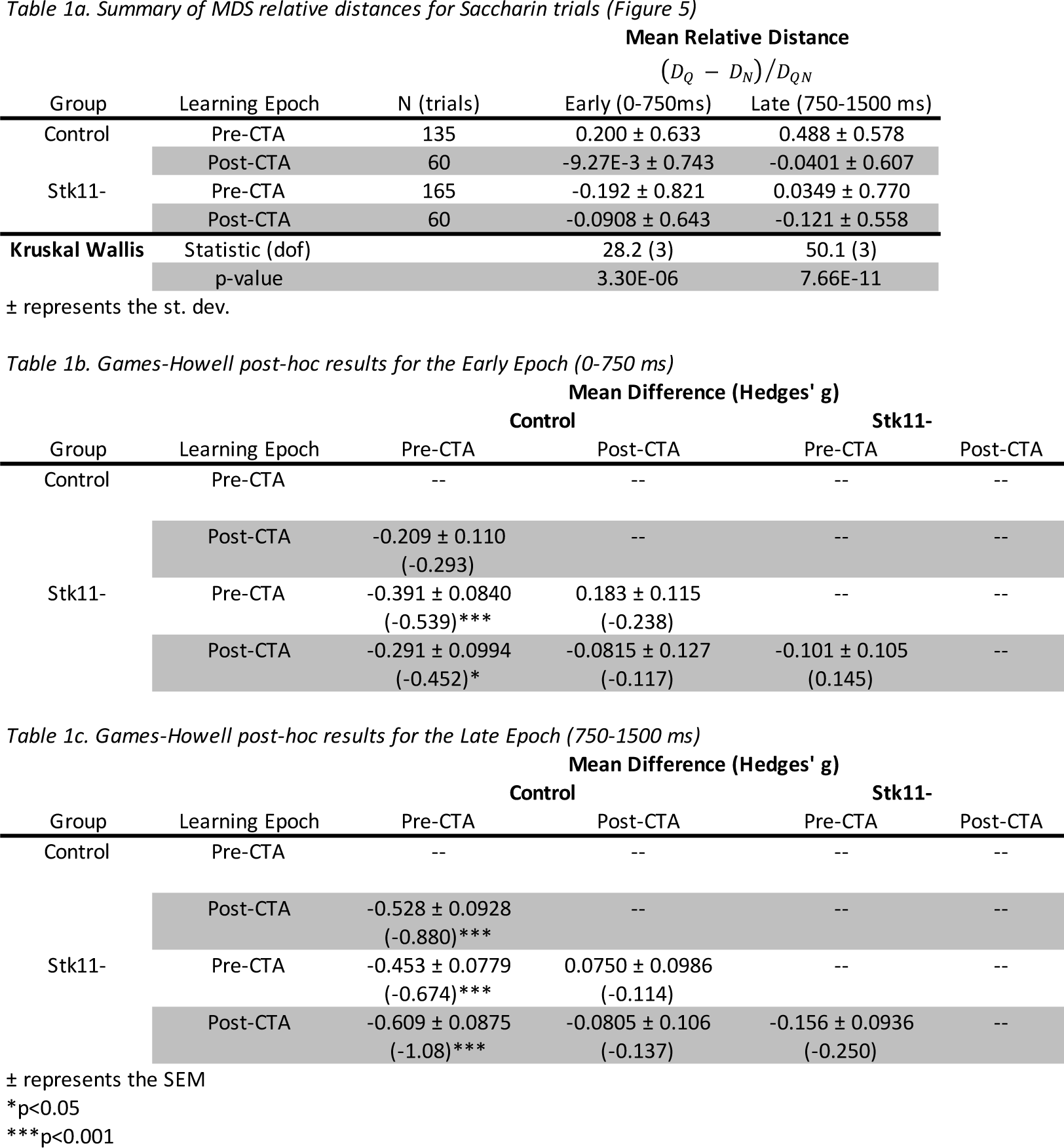
Summary of statistics for comparison of MDS relative distances presented in Figure 5.

**Figure 5.**
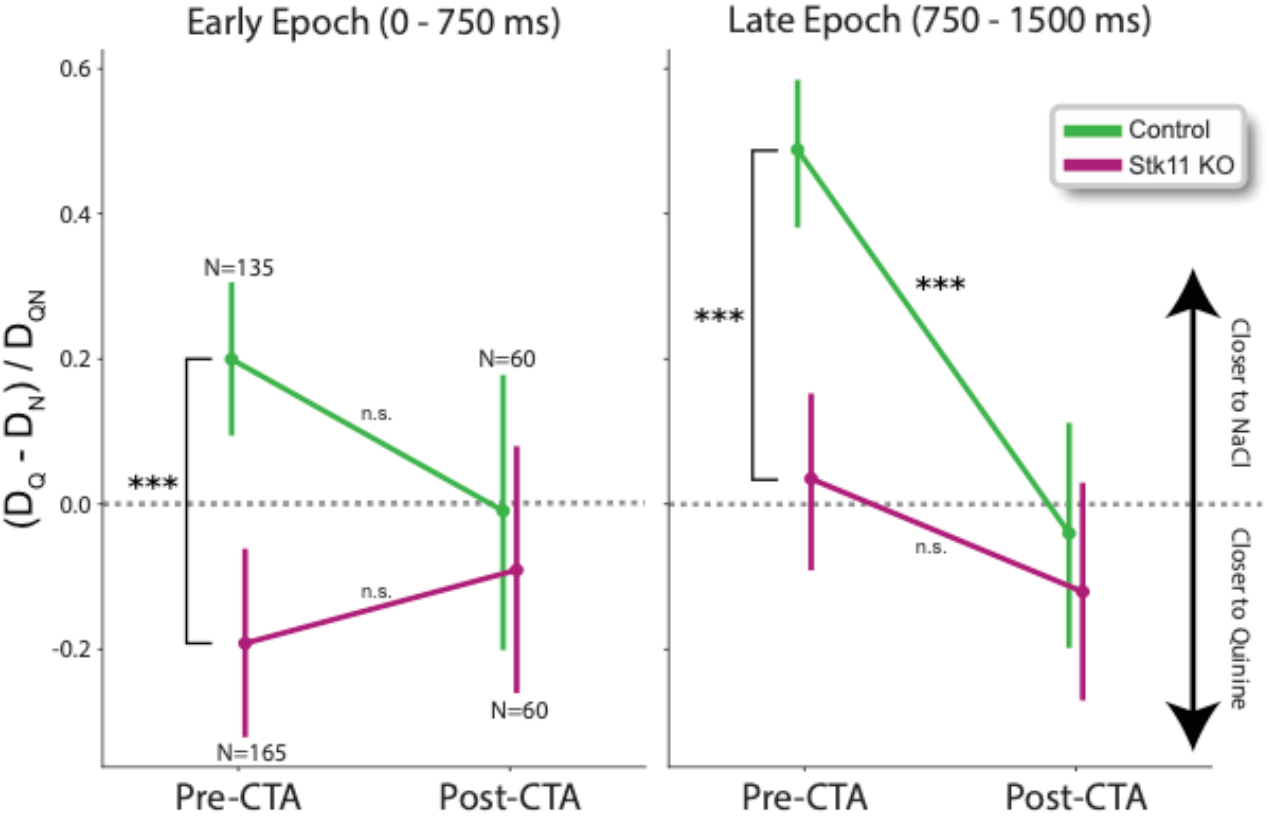
*Ensemble taste representations of saccharin are impaired* de novo *and fail to change with CTA training.* A plot of a metric showing the relative normalized distances between the saccharin representation and the average NaCl and quinine representations. Control animals showed a significant shift in the saccharin representation towards greater similarity to quinine over CTA learning in the late half of the response. Conversely, in Stk11 KO animals pre-CTA representations of saccharin did not move significantly across training. Groups were compared with non-parametric Kruskal-Wallis tests (see Table 1). Error bar reflect the 95% CI of the mean, and significance stars represent results of post-hoc Games-Howell tests; ***p<0.001.

In summary, Stk11 KO seems to disrupt the relationships among taste representations in naïve mice; furthermore, while conditioning-related response changes occur, these changes don’t impact this organization in any manner that would be expected to support learning.

### Single-trial response dynamics are altered with Stk11 KO

The data presented thus far are consistent with prior work from the Katz lab regarding the sequential processing of taste identity and palatability in GC. That prior work has also shown that palatability-related “states” arise suddenly in GC within single trials; state-to-state transition times vary widely between trials, but the overall sequence is highly reliable (Miller & Katz, 2010; Moran & Katz, 2014; Sadacca et al., 2016). CTA learning impacts the timing of these transitions (at least in rats), without altering the general dynamical structure of the responses (Moran & Katz, 2014).

To investigate the impact of Stk11 KO on these ensemble properties of GC taste responses before and after CTA training, we considered taste-evoked ensemble spike trains as multivariate Poisson processes segmented into discriminable states that could be modelled as a first-order Markov chain – that is, we applied Hidden Markov Models (HMMs) to whole ensemble responses for each taste in each session, in order to capture the underlying state dynamics of the evoked responses. Based on previous work (see also Methods) we fit 3-state models connected in a feed-forward fashion; these models account for the presence of a background state and distinct early and late response states (Figure 6a).

**Figure 6.**
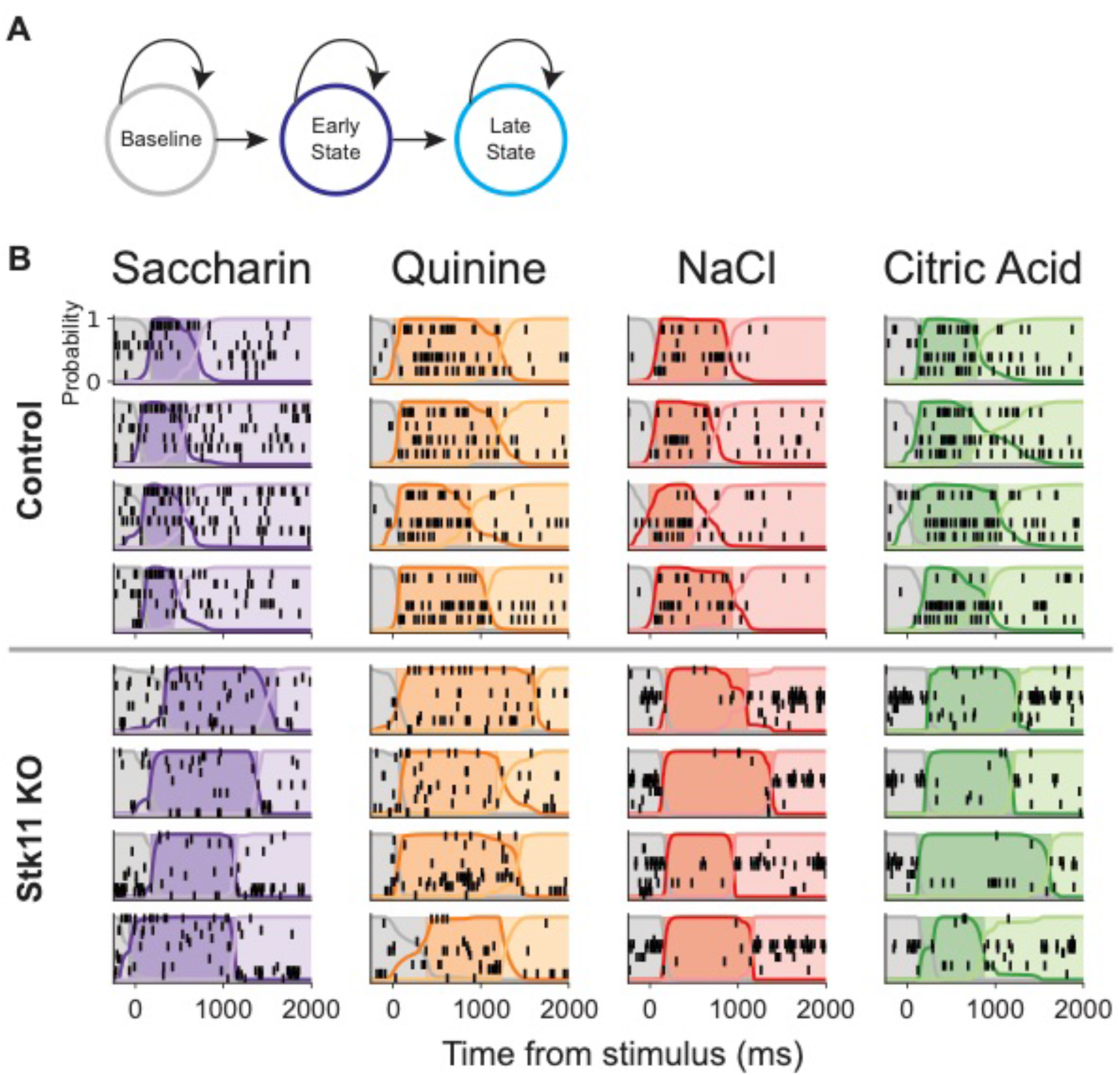
Fits of 3-state feed-forward Hidden Markov Models to GC ensemble taste responses. (A) A depiction of the hidden state sequence used to fit ensemble taste responses, with black arrows indicating possible state transitions. (B) Representative trials with HMM solutions for each tastant. Each raster shows ensemble spiking activity (black, one row per neuron) for a single trial in response to a taste delivered at time 0. The colored lines indicate the probability of being in each state at each time point (y-axis), and the background color reflects the state sequences as determined by the Viterbi decoding algorithm.

Results showed that Stk11 KO and control mice produced taste responses with similar basic state dynamics—both groups’ GC ensemble responses featured sudden and coherent firing rates changes across the population (Figure 6b). In control mice, these dynamics were largely unchanged by CTA training, a fact that represents a basic difference between mice and rats: in rats, the transition to the late (palatability) state shifts earlier in the trial with CTA training (Moran & Katz, 2014); in our control mice, this trend was at most subtle, and failed to reach significance (Figure 7A left, 7B left, 7C).

**Figure 7.**
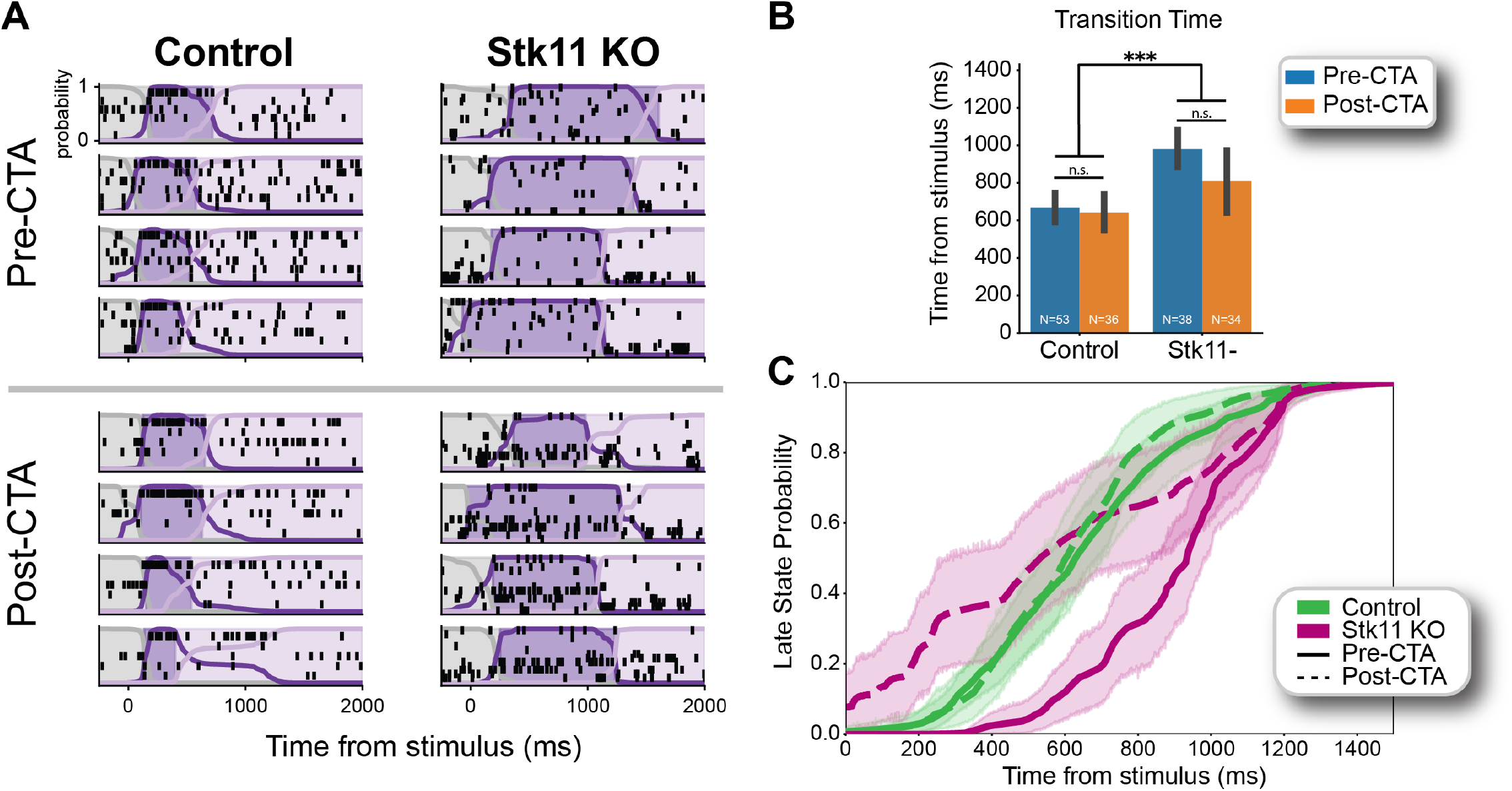
In Stk11 KO mice, state transitions are later Pre-CTA, and more variable Post-CTA. (A) Representative HMM fits for training (top) and testing (bottom) session responses to saccharin. (B) Comparison of the mean transition times from the early to the late states. Transition times were significantly later in Stk11 KO mice than in control before CTA training. Error bars represent the 95% CI of the mean. (C) Mean probability of being in the “late” state at each post-stimulus time point for each group. In Stk11 KO mice, the rise in probability pre-CTA occurred noticeably late, while in “post-CTA” testing sessions it is much more gradual (compared to the more sudden transitions seen in other groups), a result suggesting greater variability in transition times on individual trials. Shaded region depicts the 95% CI of the mean.; ***p<0.001.

More pertinent to the current hypotheses, our analysis revealed multiple differences between control and Stk11 KO transition times. For one thing, transitions were substantially delayed in naïve Stik11 KO mice: before training, this transition occurred >300ms later in the KO than control mice (Figure 7A right, 7C); a two-way ANOVA comparing the effects of group and training showed this difference between groups to be significant (F(1,157)=20, p=1.4e-5; Figure 7B).

While the average transition time for the KO moves earlier with training (by 170ms on average), no training (F(1,157)=2.8, p=0.10) or interaction effects (F(1,157)=1.7, p=0.19) were seen. This failure to identify a significant training effect, together with the fact that training also appeared to decrease the sharpness of that transition (Figure 7C magenta), led us to more closely examine the entire distribution of single-trial transition times in both groups (Figure 8a). Scrutiny of these data suggests that the post-training behavior of Stk11 KO ensembles is far from normal; further analysis (Figure 8b) reveals that whereas the distribution of transition times into the late (palatability) state are reasonably well fit by a gaussian (pre-CTA Control r^2^=0.90; post-CTA Control r^2^=0.83; pre-CTA Stk11 KO r^2^=0.74) in typical sets of saccharin responses, this structure breaks down in the Stk11 KO mice after training—the distribution loses its single strong peak (r^2^=0.040), becoming significantly different from before training (χ ^2^(1)= 41.0, p=1.5e-5, V=0.76). Clearly, the knockout of Stk11 in BLA projection neurons did not render the mice incapable of response plasticity, but this response plasticity was highly abnormal, and resulted in taste responses that were highly abnormal.

**Figure 8.**
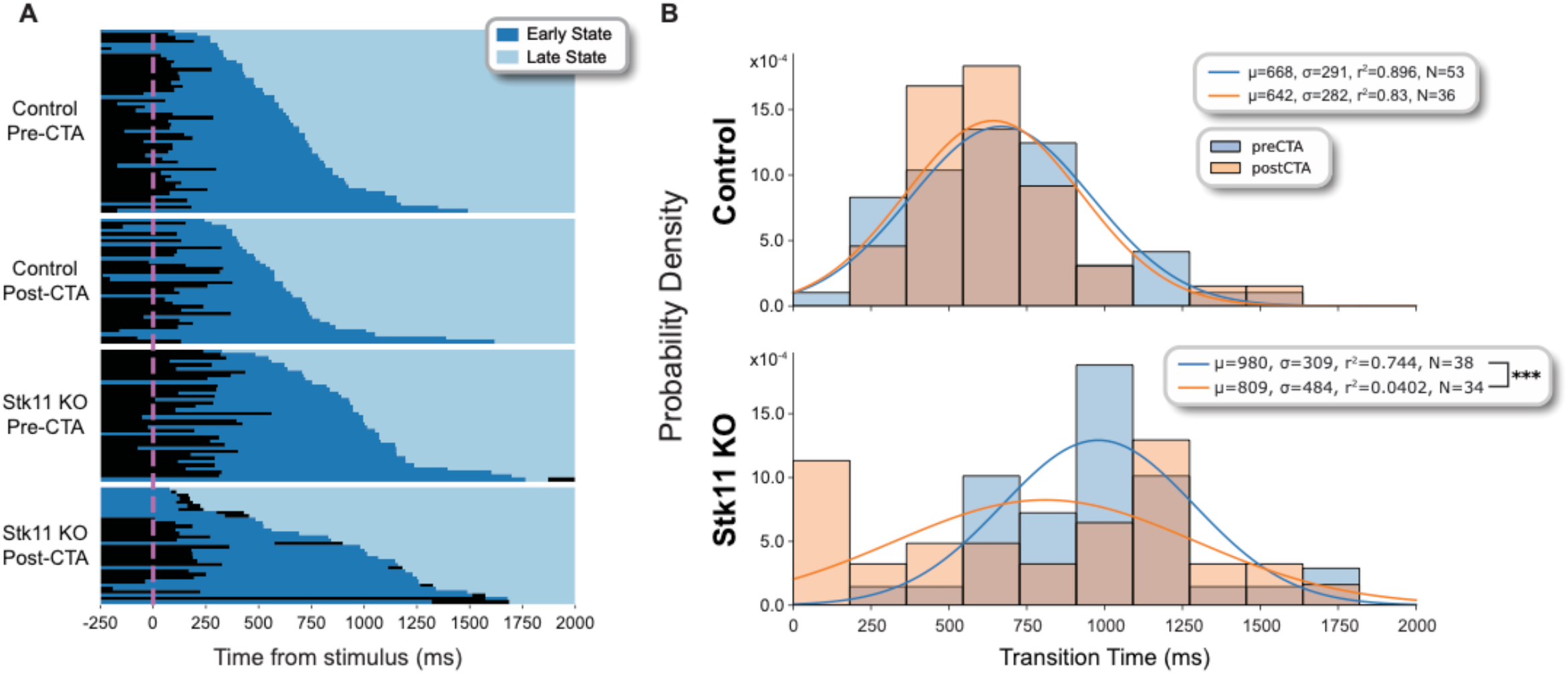
State transitions become less consistent in Stk11 KO mice following (unsuccessful) CTA training. (A) A depiction of decoded state sequences for all well-fit saccharin trials, sorted by late state onset time. Sequences produced by Stk11 KO mice show more varied structure (i.e., more gradual slope) after training. (B) Probability density histograms of transition times with lines representing gaussian fits of the distributions. χ^2^-tests reveal significantly different training/testing distributions of transition times for Stk11 KO mice (χ^2^(1)=41.0, p=1.47e-5, w=0.755) but not controls (χ^2^(1)=11.6, p=0.505, w=0.361). Furthermore, the distribution of Stk11 KO post-CTA transitions times was more poorly fit by a gaussian (r^2^=0.0402) than were all other groups (r^2^>0.7) demonstrating not just a shift in the distribution, but a loss of consistent structure in the taste responses. Significance stars represent the comparison between distributions, not between fitted lines; ***p<0.001.

## Discussion

### Mouse CTA resembles rat CTA

The gustatory neuroaxis consists of multiple brain regions acting in concert to produce precisely timed neural codes that in GC indicate, in turn, the identity and hedonic value of a tastant—information that is used to generate an appropriate behavioral response. This system supports learning whereby a tastant is associated with negative gastric consequences occurring minutes or even hours after a single ingestion session; this learning mechanism allows animals to avoid ingesting substances that have proven harmful, and is thereby evolutionarily adaptive. Here we take a step toward understanding how interactions between brain regions contribute to the formation of this robust form of learning by looking into how the knockout of a single gene in BLA projection neurons impacts ensemble activity in GC during taste processing and over the course of CTA training.

Prior studies using rats have shown that it is primarily the late “epoch” of the taste response that changes over the course of CTA training, as the code continues to reflect the (changing) value of the tastant and reliably predict behavior (Grossman et al., 2008; Sadacca et al., 2016; Mukherjee et al., 2019). This remains true in this study – in control mice (that did learn the CTA), changes in single unit responses to saccharin primarily occurred in the late epoch (>750ms) with CTA training (Figure 4), such that the ensemble representations of saccharin during this epoch shifted dramatically from resembling another palatable taste (NaCl) towards more resembling an unpalatable taste (quinine) over learning (Figure 5). In fact, the only rat result that we were unable to clearly replicate in control mice was a significant decrease in the onset latency of the late state over CTA learning (Moran & Katz, 2014); a trend toward this shift was observed (Figure 7B&C, Figure 8B), but the magnitude was smaller (in our hands insignificantly so) than that seen in rats.

It’s worth noting that in our control mice, learned aversions to saccharin were mild—a fact that might have made it difficult to observe a reduction in transition latency. This is probably a reflection of the fact that: 1) we tested learning only after first performing post-training electrophysiological recordings during which mice received saccharin without any associated adverse effects; and 2) we tested responses to a whole battery of tastes both just before and just after the training sessions. Both of these features of our preparation would be likely to interfere with the expression of learning—by the time we assessed CTA, some extinction (Nolan et al., 1997; Bouton, 2004; Moran & Katz, 2014) had probably occurred because of the safe presentation of saccharin, and some diffusion (Richardson et al., 1984; McLaren & Mackintosh, 2002; Heyer et al., 2003) or overshadowing (Hall & Symonds, 2006; Kwok et al., 2012) of learning had probably also occurred because of the peri-training presentation of other tastes.

### Stk11KO impacts taste processing

While our prior work (Levitan et al., 2020) did not unearth significant differences in the overall structure of taste processing caused by Stk11 KO, the more in-depth analysis performed here reveals some interpretable abnormalities in the neural taste processing of mice lacking Stk11 in BLApn. At the most basic level, putative pyramidal neurons in GC show slightly higher basal firing rates in Stk11 KO than control mice—a change easily attributable to the previously-noted increase of excitability in BLApn, and to the subsequent increase of excitatory drive into GC from BLA. Together with the observed elevation in palatability-relatedness of firing throughout the taste response in Stk11 KO animals (Figure 3), these data suggest that one result of losing BLApn Stk11 is an alteration in the impact of BLA on GC, such that the magnitude of the transfer of information from one to the other is enhanced.

This latter novel result may have been previously undetectable due to our prior study’s inclusion of sucrose in the taste battery: as sucrose has reliably high palatability (and correspondingly large differences in response firing rates from other tastants), it is likely that its inclusion swamps the impact of other (subtler) response differences on the correlation computation. By only considering the responses of the NaCl, quinine and citric acid here, we may actually obtain a clearer picture of how these subtler taste response differences lay out with respect to taste palatability. This conclusion is consistent with the suggestion that the loss of Stk11 enhances the contribution of BLA to GC activity (*via* increases in PN excitability).

We also observed a difference in how naïve Stk11 KO mice process saccharin. Typically, saccharin taste responses are more similar to NaCl than to quinine, by virtue of the palatability-relatedness of late epoch coding and the assumption that NaCl and saccharin are similarly palatable (Moran & Katz, 2014). In Stk11 KO mice, however, the *de novo* saccharin representation is significantly closer to quinine than controls in both epochs of the taste response. Though separable from the actual calculation of palatability (which was evaluated in relation to coding of quinine, NaCl, and citric acid), this does suggest a certain corruption of information content – or the organization of information – in the GC of Stk11 KO mice. Future work will investigate whether perhaps Stk11 KO mice naturally find saccharin less appealing than do control mice, as has been found with other mouse strains and manipulations (Yirmiya et al., 1988; Harkin et al., 2002; O’Brien et al., 2013; Awad et al., 2020).

Finally, the ensemble dynamics of naïve taste processing are also altered by KO of Stk11 in BLApn, in that the onset of the late state in single trials tends to be greatly delayed prior to training (Figure 7B). With this transition being directly linked to the behavioral decision of whether to spit or swallow the tastant (Sadacca et al., 2016; Mukherjee et al., 2019), this large delay in the occurrence of this transition merits further investigation as it could mean a large delay in making a potentially life-saving decision; alternatively, it could mean that these abnormally timed and coded states no longer command the same behavioral relevance in the KO mice. Further experiments looking at both amygdalar responses to tastants and the timings of orofacial responses to tastants will place the changes in GC taste processing observed here into richer context, and thereby help to explain further how the circuit is disrupted.

### CTA-related response plasticity is abnormal (but present) in Stk11KO mice

Our naïve prediction was that the lack of learning in Stk11KO mice should be reflected in a concomitant lack of response plasticity. There was ample *prima facie* reason to doubt this simplest hypothesis, however, starting with the fact that the impact of a loss of Stk11 in BLApn is actually to increase these neurons’ basal excitability (Levitan et al., 2020) – a change conceptually associated with increases in network plasticity (Cohen et al., 1999; Malik & Chattarji, 2012; Yiu et al., 2014).

In fact, the initial hypothesis proved premature: we observed that GC saccharin responses of single neurons in Stk11 KO animals did change significantly with CTA training (Figure 4); these changes were not “learning-appropriate,” however, in that they centered on a period prior to the palatability epoch (i.e., <750ms post-stimulus delivery) that is involved in stimulus identification (Yoshida & Katz, 2011), and left the later, normally learning-sensitive phase of the responses, relatively untouched. In fact, the later onset of the palatability state in these mice (at ∼980ms post-stimulus delivery) places the training-related changes seen firmly within this earlier identification epoch of the taste response. Furthermore, the GC coding dynamics in Stk11 KO mice, which (again) are already abnormal prior to learning, are rendered incoherent by CTA training, with the very organization of state-to-state transition times falling apart (Figure 8B, bottom). In summary, the course of CTA training causes plastic changes in mice lacking Stk11 in BLApn, but the nature of the changes is such that they disrupt the processing of the training taste and fail to support the learned behavior.

If anything, the changes in Stk11 KO taste responses wrought by our training regimen are specifically unsupportive of CTA. While single neurons exhibit earlier changes in their responses to saccharin, the representational content (i.e., the relationship between saccharin responses and those to tastes whose positive or negative hedonics are untouched by training) remains mostly constant. Not only does “learning” impact the “wrong” portion of the response, that influence fails to change coding in the right direction. These data further suggest that the loss of Stk11 alters the timing of the impact of BLA on GC, changing the window of the taste response impacted by the training.

Finally, it is important to consider the fact that GC, like most sensory cortices (Olshausen & Field, 2004; Niell & Stryker, 2008; Poo & Isaacson, 2009; Sakata & Harris, 2009; Crochet et al., 2011), normally relies on sparse firing to encode and process gustatory information. The increased input from BLA results in increased activity in GC; baseline firing rates (in putative pyramidal neurons) are higher, more neurons are active in response to tastes appearing on the tongue, and palatability-related firing is enhanced. A consequence of this abnormality is the augmentation of GC’s normal sparse coding scheme with additional (but not necessarily useful) information. It is entirely possible, given this fact, that the loss of Stk11 in BLApn alters the perceptual properties of specific tastants (our data in fact suggests that saccharin might be perceived differently by KO mice). If so, even if naïve mice can distinguish tastes, the “trained” mouse might fail to identify the tastant associated with the gastric malaise as the same tastant presented prior to training. This abnormality is also consistent with the disturbed organization of taste representations seen in Stk11 KO mice prior to – and over the course of – CTA training (Figure 5). In future work, we will investigate whether Stk11 KO mice can discriminate between tastants reliably, and whether they can identify a conditioned tastant as the same stimulus that had been presented prior to CTA training.

## Materials and Methods

### Subjects

Male Stk11^fl/fl^ (B6(Cg)^Stk11tm1.1Sjm^/J) mice aged P60-P90 were used for these experiments. Mice were kept on a 12-hour light-dark cycle and given ad libitum access to food and water except during experiments habituation and recording days, at which time water access was restricted, while food remained available ad libitum. All procedures were approved by the Brandeis University Institutional Animal Care and use Committee (IACUC) in accordance with NIH guidelines.

### Surgery

All animals were habituated to the testing environment prior to surgery. Mice were initially anesthetized with an I.P. injection of a ketamine/xylazine (K.X.) mixture (50 mg/kg & 2.66 mg/kg, respectively). Their heads were shaved and then inserted into the stereotaxic frame (David Kopf Instruments; Tujunga, CA). Anesthesia was maintained with 0.5 – 2% isoflurane. The skull was exposed and cleaned, and craniotomies were made for BLA virus injection (−1.4 mm AP, ±3.4 mm ML) and GC electrode implantation (+1.0mm AP, ±3.4mm ML). BLA was injected bilaterally with AAV2/5-Camk2α::Cre-GFP (Stk11 KO; N=6) or a control virus consisting of AAV2/5-Camk2α::GFP (Control; N=10) (UNC, vector core) using sterile glass micropipettes (10-20μm diameter) attached to a partially automated microinjection device (Nanoject III Microinjector, Drummond Scientific; Broomall, PA). The micropipettes were slowly lowered into BLA at 4.6 mm then 4.3 mm from the dura; At each depth, 200nL of the viral solution were delivered via 10 pulses of 20nL every 10 seconds. The micropipette was retracted 10 minutes after the dorsal-most injection on each side. Small burr holes were drilled anterior to lambda on either side of the midline and M1 stainless steel screws were inserted both to act as grounding contacts and to anchor the assembly to the skull. Two 16-wire bundles of formvar-coated 25 μm nichrome wire attached to mini-microdrives were slowly lowered to 0.5mm above ventral GC (2.5 mm from the dura) as done previously (Levitan et al., 2019; Levitan et al., 2020) and secured in place with C&B Metabond (Parkell; Edgewood, NY). To aid with later identification of electrode locations, bundle tips were lightly coated with DiL stain (ThermoFisher Scientific; Indianapolis, IN) before implantation.

Mice were given a booster I.P. injection of the original induction mix 10 minutes before being briefly removed from the stereotaxic frame, so that an intraoral cannula (IOC; flexible plastic tubing) could be inserted close to the tongue inside the cheek (Fontanini & Katz, 2006), after which they were replaced in the stereotaxic frame, and stainless-steel grounding wire was connected from the circuit board to the grounding screws. The craniotomies were sealed with Kwik-Sil (World Precision Instruments; Sarasota, FL) and components were embedded in dental acrylic and thereby secured to the skull.

Mice were allowed to recover on a heating mat before being returned to their home cages. Post-operative injections (s.c.) of an analgesic (Metacam; 2 mg/kg; Boehringer Ingelheim Animal Health USA Inc.; Duluth, GA), lactated ringer’s solution (0.4 ml) and an antibiotic (Pro-Pen-G; 150,000 U/kg; Bimeda; Oakbrook Terrace, IL) were given immediately following surgery and at 24hr and 48hr later. Injections of lactated ringer’s solution continued for several extra days until weight stabilized.

### Behavioral Procedures

After 7d of recovery from surgery, mice were habituated to the testing environment for 30 min per day. At post-op day 10, animals were placed on a water restriction schedule for the remainder of the experiment (6 days) and habituated to receiving ∼1ml (60 pulses, 16-18 μL per pulse, 15-25ms between pulses) of water through the IOC in the morning (they also received 10 minutes of free water access in their home cages in evening). On Post-op days 11 & 12, electrode bundles were lowered a total of 0.5mm, using the built-in microdrive assembly; recording started 24hrs later—4 sessions over 3 days. The first session (Tastes 1) consisted of 60 IOC deliveries of a battery of 4 taste solutions (15 deliveries per taste, ∼16 μL per delivery, 15-25ms between deliveries). These solutions were water, 0.02M citric acid, 0.1M NaCl and 0.001M quinine (concentrations chosen to maximize comparability with earlier papers (Moran & Katz, 2014; Levitan et al., 2019)); 2 tastes were delivered in pseudo-random order for the first half of the session, and then the others. Pairings differed for each animal, but citric acid and quinine were never paired, in order to prevent animals from receiving too many consecutive un-palatable taste deliveries (Tastes 1).

The next day mice were given 60 deliveries of saccharin (0.02 M, 16-18μL per delivery; CTA Train session). Due to an equipment malfunction, 2 animals were only given 45 deliveries (20-21μL per delivery) to keep the amount of total amount of saccharin consumed during training the same across animals. Five minutes after the final saccharin delivery, they were given an I.P. injection of 0.15M LiCl (2% b.w.) to induce gastric malaise. This procedure allows, under normal circumstances, a selective association (condition taste aversion, CTA) to form between the malaise and the novel tastant. One day later, animals received 15 deliveries of saccharin (CTA Test), and 1-2hr later underwent another session of the battery of 4 tastes (Tastes 2). The evening after the final recording session, instead of 10 min access to water, animals received 10 min access to saccharin solution in order to evaluate strength of CTA. To assess CTA, the amount of saccharin consumed was compared to water consumption on previous nights. Based on prior work in the lab (and since the earlier consumption of saccharin as well as other tastes likely resulted in a certain mildness of learning), we chose a saccharin consumption of 80% of average nightly water consumption as a conservative threshold, below which animals were deemed to have learned the CTA; this was an adequate criterion for our purposes because the primary measure was difference between groups, and thus the prediction was that many control mice would reach this criterion but that few Stk11KO mice would do so. 5 of the 10 control mice reached this learning criterion, and 5 of the 6 Stk11 KO mice did not. The one Stk11 KO mouse whose saccharin consumption was below the threshold was excluded from analysis because that animal’s nightly water consumption steadily decreased over habituation days, and we could not be sure if its lack of saccharin consumption was an effect of CTA learning or some other impairment resulting in reduced intake.

### Electrophysiology

During taste presentations, up to 32 extracellular potentials were simultaneously sampled and digitized by an Intan RHD2132 headstage and sent to a DAQ @ 30kHz (Intan Technologies; Los Angeles, CA). These time series, as well as time-stamped records of stimulus onset and offset times, were stored for offline analysis in python using the blechpy analysis package: discriminable action potentials with at least a 3:1 signal-to-noise ratio were isolated offline by amplitude thresholding; 2ms snippets of each waveform were extracted, up-sampled 10x and aligned to the wave peak. The UMAP algorithm was used to extract waveform features, which were fit by Gaussian mixture models to cluster spiking events—clusters that were then manually curated (by experimenters blind to taste responses and injection type) to ensure conservatism. Only clusters clearly separated in UMAP or PCA space, and containing no 1ms ISI violations and fewer than 1% 2ms ISI violations, were deemed to reflect single-neuron activity. These methods and criteria are in accordance with previously published methods from our lab (Moran & Katz, 2014; Sadacca et al., 2016; Levitan et al., 2019; Mukherjee et al., 2019).

### Histology

Animals were deeply anesthetized with isoflurane and euthanized with a K.X. overdose. Animals were then perfused transcardially with 0.9% saline solution followed by 4% formaldehyde in PBS. Brains were extracted and stored in 4% formaldehyde, 30% sucrose solution. Once fully fixed, 50 μm coronal sections of BLA and GC were visualized under a fluorescent microscope (KEYENCE BZ-X800; Itasca, IL) to verify injection and electrode positions.

### Detection of single neurons reliably held across sessions

To determine if a waveform recorded from a single electrode for multiple sessions represented the same single neuron, we used a spike shape analysis successfully used in several previous studies (Nicolelis et al., 2003; Grossman et al., 2008; Herry et al., 2008; Moran & Katz, 2014). A non-parametric clustering statistic (J_3_) was used to quantify the separation and grouping between 2 clusters. J_3_-values were computed for all single neurons across all sessions using spikes from the first and last thirds of each recording session to generate a distribution of intra-session J_3_ values of known single neurons. Then, for each pair of single cells on the same electrode between 2 consecutive sessions, the J_3_ was computed and compared to the intra-session distribution. A conservative criterion was applied: only pairs of units with an inter-session J_3_ value less than the 95^th^ percentile of the intra-session J_3_ distribution (for the entire experimental group) were deemed to be held across sessions. This mean that only pairs of units from across days that are more similar to each other than at least 5% of single units are to themselves within a recording session.

### Analysis of single neuron taste responses

Single neurons were classified as regular spiking or fast-spiking based on inspection of spike waveform shape and ISI histograms (for details see Levitan et al., 2019). Regular spiking cells were taken as putative pyramidal neurons while fast-spiking cells were labelled as interneurons. Baseline firing rates were computed as the average spike rates over the 1s period prior to stimulus deliveries. Response firing rates were initially computed to be the average spike rate over the entire 1.5s post-stimulus period. Two-way ANOVAs were used to assess the effects of group (Stk11 KO vs Control) and training (pre- vs post-CTA) on baseline and response firing rates for each unit type (Figure 2).

Using the data from the pre- and post-CTA taste battery sessions (Tastes 1 and Tastes 2), we also looked at the correlation between the response firing rates and the rank-order hedonic values of the given tastants (Quinine [1], Citric Acid [2] and NaCl [3]; Figure 3). This allowed us to compare the decision-relevant information content of the single-neuron responses between groups. Firing rates were computed in non-overlapping 250ms bins. Within each time bin, the Spearman correlation coefficient (R^2^) was computed for the correlation between the ranked firing rates and the palatability values. A two-way ANOVA was used to investigate the effects of group and training on the correlation of responses to palatability. All ANOVA analyses used Bonferroni corrected post-hoc t-tests for within group comparisons.

### Changes in held neuron responses to Saccharin over training

To understand the time course of changes to single neuron responses to Saccharin over training, we analyzed the responses of neurons that were deemed reliably held from the CTA Train session to the CTA Test session (Figure 4). For these, firing rates were computed in six consecutive, non-overlapping 250ms bins post-stimulus, similar to the manner used previously (Grossman et al., 2008). Response changes in each time bin were evaluated using a one-way ANOVA (p-values were Bonferroni corrected), and the number of neurons showing significant changes across training in each time bin was compared between groups using a χ^2^-test.

Due to our conservative criteria for held neurons, this analysis was the limiting factor for determining sample size needed for our study. Preliminary data suggested this analysis would have an effect size of around 0.960 (Cramer’s V), which would have required a total sample of 15 neurons to reliably detect at a power 0.95 (computed using G*Power 3.1; see Faul et al., 2009). Post-hoc analysis found that our final effect size was 0.580; with a final sample size of 38 held cells, this resulted in a final power of 0.946 for our analysis.

### Multidimensional Scaling (MDS)

Multidimensional scaling allows for the reduction of dimensionality in multi-variate data while preserving the relative two-dimensional distances between points. We used MDS, as has been done many times previously (e.g. Chang & Scott, 1984; Geran & Travers, 2006; Moran & Katz, 2014), to analyze the similarity between saccharin taste responses and those of NaCl (palatable) and quinine (unpalatable). Activity projections into 2D space were computed separately for the early (0-750ms) and late (750-1500ms) epochs of the response; in each, Euclidean distances were computed between representations of individual saccharin responses and the mean quinine (D_Q_) and NaCl representations (D_N_) for each recording set. These distances were then normalized by the distance between the NaCl and quinine means (D_QN_) to provide a dimensionless distance metric

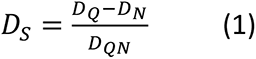

which quantifies the similarity of the saccharin response to the NaCl and quinine representations: positive values indicate that the saccharin representation is more similar to NaCl than quinine, while negative values indicate the opposite (Figure 5). The populations for this analysis consisted only of units that were reliably held from Tastes 1 to CTA Train (pre-CTA) or from CTA Test to Tastes 2 (post-CTA), and any recording sets that did not have at least 2 held units across these periods were excluded from this analysis.

### Poisson Hidden Markov Models (HMMs)

To ascertain the underlying temporal structure of single trial ensemble responses, we fit sets of simultaneously-recorded spike trains to Poisson HMMs (Rabiner, 1989; Abeles et al., 1995; Jones et al., 2007; Kemere et al., 2008; Miller & Katz, 2010; Moran & Katz, 2014; Sadacca et al., 2016). Since our goal was to look at the timing of these state transitions (and not to have the model classify taste responses), we fit separate models for each tastant for each recording session. We constrained the search space for our parameters, fitting models for −250 ms to 2000 ms peri-stimulus with 2-6 states connected in a feed-forward manner. The Bayesian Information Criteria (BIC) was used to evaluate model fit, and while we saw no statistically significant differences in BIC relative to the number of states, 2 or 3 state models consistently fit the data best (i.e., had the lowest BIC values). Given the extensive evidence (Katz et al., 2001; Fontanini & Katz, 2006; Miller & Katz, 2010; Piette et al., 2012; Sadacca et al., 2012; Moran & Katz, 2014; Sadacca et al., 2016; Levitan et al., 2019) that cortical taste responses consist of 2 major chemosensory coding states or epochs, we settled on models containing 3 states connected in feed-forward manner (Figure 6a) so as to account for the presence of a background state, an early state and a late state.

Spike trains were provided to the model at 1ms resolution, and initial model firing rates were randomly selected from a normal distribution using the mean and standard deviation for each units’ firing rates across trials. Values for the initial transition matrix were drawn randomly from a normal distribution with mean 0.05 and standard deviation of 0.01. Diagonal values (the probability of remaining in the same state at each time step) were similarly drawn from a normal distribution with mean of 0.99 and sigma of 0.01 to ensure a high probability of remaining in the same state, and the elements corresponding to transitions backward or advancing more than 1 state were set to zero. Finally, all rows were normalized to sum to 1.

For each dataset, models were initialized and fit 50 separate times; in each run, fitting terminated when the change in log likelihood over iterations dropped below 1e-10. The model with the highest log likelihood was deemed the best description of the data. Fitting was done using the Baum-Welch algorithm (Rabiner, 1989), which provided the probabilities of the ensemble being in each state at each time point for each trial (Figure 7c). The optimal sequences of states for each trial were decoded using the Viterbi decoding algorithm (Viterbi, 1967), which provides the maximum likelihood decoding of each trial based on the fitted model. While the 3-state models routinely provided good fits to the data (Figure 6b), for subsequent analysis we nonetheless chose early and late states independently, designating them to be the most probable sequence of states present in the bins 200-700ms and 750-1500ms respectively.

Only recording sessions containing at least 3 well-isolated single cells were included in any such analysis, to avoid allowing results to be unduly biased toward the dynamics of single neurons. Furthermore, only the first 15 saccharin trials of CTA-Train sessions were used for this analysis since all other sessions only had 15 trials per tastant to fit; this ensured that model quality and hyperparameters (number of states, time window, bin size, etc.) could remain constant across all trained models.

### Analysis of HMM Transition Times

As the offset times for early states and onset times for the late states were reliably coupled, we defined transition times as the offset times for the early states. Mean transition times (Figure 7B) were compared using a two-way ANOVA for group (Stk11 KO vs Control) and training (pre- vs post-CTA) effects. However, as inspection of the distributions revealed large variability in the transition times of Stk11 KO responses post-CTA, these distributions were then compared between pre- and post-CTA within each group using χ^2^-tests, and fit to Gaussian functions (Figure 8B) to quantify the differences in transition distributions. Only trials decoded as having both the early and late states present post-stimulus were used in these analyses.

## Acknowledgements

This research was supported by US National Institutes of Health grant R21-MH104318 and the National Institute on Deafness and Other Communication Disorders (NIDCD) grants DC006666 and DC007703.

## References

Abeles, M., Bergman, H., Gat, I., Meilijson, I., Seidemann, E., Tishby, N., & Vaadia, E. (1995). Cortical activity flips among quasi-stationary states. Proc Natl Acad Sci U S A, 92(19), 8616–8620. doi:10.1073/pnas.92.19.8616

Arieli, E., Gerbi, R., Shein-Idelson, M., & Moran, A. (2020). Temporally-precise basolateral amygdala activation is required for the formation of taste memories in gustatory cortex. J Physiol, 598(23), 5505–5522. doi:10.1113/JP280213

Awad, G., Roeckel, L. A., Massotte, D., Olmstead, M. C., & Befort, K. (2020). Deletion of mu opioid receptors reduces palatable solution intake in a mouse model of binge eating. Behav Pharmacol, 31(2&3), 249–255. doi:10.1097/FBP.0000000000000496

Beirowski, B. (2019). The LKB1-AMPK and mTORC1 Metabolic Signaling Networks in Schwann Cells Control Axon Integrity and Myelination: Assembling and upholding nerves by metabolic signaling in Schwann cells. Bioessays, 41(1), e1800075. doi:10.1002/bies.201800075

Bouton, M. E. (2004). Context and behavioral processes in extinction. Learning & memory (Cold Spring Harbor, N.Y.), 11(5), 485–494. doi:10.1101/lm.78804

Bures, J., Bermudez-Rattoni, F., & Yamamoto, T. (1998). Conditioned Taste Aversion: Memory of a Special Kind: Oxford University Press.

Chang, F. C., & Scott, T. R. (1984). Conditioned taste aversions modify neural responses in the rat nucleus tractus solitarius. The Journal of Neuroscience, 4(7), 1850–1862. doi:10.1523/jneurosci.04-07-01850.1984

Cohen, A. S., Coussens, C. M., Raymond, C. R., & Abraham, W. C. (1999). Long-lasting increase in cellular excitability associated with the priming of LTP induction in rat hippocampus. J Neurophysiol, 82(6), 3139–3148. doi:10.1152/jn.1999.82.6.3139

Crochet, S., Poulet, J. F., Kremer, Y., & Petersen, C. C. (2011). Synaptic mechanisms underlying sparse coding of active touch. Neuron, 69(6), 1160–1175. doi:10.1016/j.neuron.2011.02.022

Faul, F., Erdfelder, E., Buchner, A., & Lang, A. G. (2009). Statistical power analyses using G*Power 3.1: tests for correlation and regression analyses. Behav Res Methods, 41(4), 1149–1160. doi:10.3758/BRM.41.4.1149

Fonseca, E., Sandoval-Herrera, V., Simon, S. A., & Gutierrez, R. (2020). Behavioral Disassociation of Perceived Sweet Taste Intensity and Hedonically Positive Palatability. eNeuro, 7(5). doi:10.1523/ENEURO.0268-20.2020

Fontanini, A., & Katz, D. B. (2006). State-dependent modulation of time-varying gustatory responses. J Neurophysiol, 96(6), 3183–3193. doi:10.1152/jn.00804.2006

Gallo, M., Roldan, G., & Bures, J. (1992). Differential involvement of gustatory insular cortex and amygdala in the acquisition and retrieval of conditioned taste aversion in rats. Behav Brain Res, 52(1), 91–97. doi:10.1016/s0166-4328(05)80328-6

Garcia-Delatorre, P., Perez-Sanchez, C., Guzman-Ramos, K., & Bermudez-Rattoni, F. (2014). Role of glutamate receptors of central and basolateral amygdala nuclei on retrieval and reconsolidation of taste aversive memory. Neurobiol Learn Mem, 111, 35–40. doi:10.1016/j.nlm.2014.03.003

Geran, L. C., & Travers, S. P. (2006). Single Neurons in the Nucleus of the Solitary Tract Respond Selectively to Bitter Taste Stimuli. Journal of Neurophysiology, 96(5), 2513–2527. doi:10.1152/jn.00607.2006

Grossman, S. E., Fontanini, A., Wieskopf, J. S., & Katz, D. B. (2008). Learning-related plasticity of temporal coding in simultaneously recorded amygdala-cortical ensembles. J Neurosci, 28(11), 2864–2873. doi:10.1523/JNEUROSCI.4063-07.2008

Hall, G., & Symonds, M. (2006). Overshadowing and latent inhibition of context aversion conditioning in the rat. Auton Neurosci, 129(1-2), 42-49. doi:10.1016/j.autneu.2006.07.013

Harkin, A., Houlihan, D. D., & Kelly, J. P. (2002). Reduction in preference for saccharin by repeated unpredictable stress in mice and its prevention by imipramine. J Psychopharmacol, 16(2), 115–123. doi:10.1177/026988110201600201

Herry, C., Ciocchi, S., Senn, V., Demmou, L., Muller, C., & Luthi, A. (2008). Switching on and off fear by distinct neuronal circuits. Nature, 454(7204), 600–606. doi:10.1038/nature07166

Heyer, B. R., Taylor-Burds, C. C., Tran, L. H., & Delay, E. R. (2003). Monosodium glutamate and sweet taste: generalization of conditioned taste aversion between glutamate and sweet stimuli in rats. Chem Senses, 28(7), 631–641. doi:10.1093/chemse/bjg056

Jones, L. M., Fontanini, A., Sadacca, B. F., Miller, P., & Katz, D. B. (2007). Natural stimuli evoke dynamic sequences of states in sensory cortical ensembles. Proc Natl Acad Sci U S A, 104(47), 18772–18777. doi:10.1073/pnas.0705546104

Katz, D. B., Simon, S. A., & Nicolelis, M. A. L. (2001). Dynamic and Multimodal Responses of Gustatory Cortical Neurons in Awake Rats. The Journal of Neuroscience, 21(12), 4478–4489. doi:10.1523/jneurosci.21-12-04478.2001

Kemere, C., Santhanam, G., Yu, B. M., Afshar, A., Ryu, S. I., Meng, T. H., & Shenoy, K. V. (2008). Detecting neural-state transitions using hidden Markov models for motor cortical prostheses. J Neurophysiol, 100(4), 2441–2452. doi:10.1152/jn.00924.2007

Kwok, D. W., Livesey, E. J., & Boakes, R. A. (2012). Serial overshadowing of taste aversion learning by stimuli preceding the target taste. Learn Behav, 40(4), 427–438. doi:10.3758/s13420-011-0064-0

Lavi, K., Jacobson, G. A., Rosenblum, K., & Luthi, A. (2018). Encoding of Conditioned Taste Aversion in Cortico-Amygdala Circuits. Cell Rep, 24(2), 278–283. doi:10.1016/j.celrep.2018.06.053

Levitan, D., Lin, J. Y., Wachutka, J., Mukherjee, N., Nelson, S. B., & Katz, D. B. (2019). Single and population coding of taste in the gustatory cortex of awake mice. J Neurophysiol, 122(4), 1342–1356. doi:10.1152/jn.00357.2019

Levitan, D., Liu, C., Yang, T., Shima, Y., Lin, J. Y., Wachutka, J., … Nelson, S. B. (2020). Deletion of Stk11 and Fos in mouse BLA projection neurons alters intrinsic excitability and impairs formation of long-term aversive memory. Elife, 9. doi:10.7554/eLife.61036

Li, J. X., Maier, J. X., Reid, E. E., & Katz, D. B. (2016). Sensory Cortical Activity Is Related to the Selection of a Rhythmic Motor Action Pattern. J Neurosci, 36(20), 5596–5607. doi:10.1523/JNEUROSCI.3949-15.2016

Lin, J. Y., Mukherjee, N., Bernstein, M. J., & Katz, D. B. (2021). Perturbation of amygdala-cortical projections reduces ensemble coherence of palatability coding in gustatory cortex. Elife, 10. doi:10.7554/eLife.65766

Maffei, A., Haley, M., & Fontanini, A. (2012). Neural processing of gustatory information in insular circuits. Curr Opin Neurobiol, 22(4), 709–716. doi:10.1016/j.conb.2012.04.001

Malik, R., & Chattarji, S. (2012). Enhanced intrinsic excitability and EPSP-spike coupling accompany enriched environment-induced facilitation of LTP in hippocampal CA1 pyramidal neurons. J Neurophysiol, 107(5), 1366–1378. doi:10.1152/jn.01009.2011

McLaren, I. P., & Mackintosh, N. J. (2002). Associative learning and elemental representation: II. Generalization and discrimination. Anim Learn Behav, 30(3), 177–200. doi:10.3758/bf03192828

Miller, P., & Katz, D. B. (2010). Stochastic transitions between neural states in taste processing and decision-making. J Neurosci, 30(7), 2559–2570. doi:10.1523/JNEUROSCI.3047-09.2010

Moran, A., & Katz, D. B. (2014). Sensory cortical population dynamics uniquely track behavior across learning and extinction. J Neurosci, 34(4), 1248–1257. doi:10.1523/JNEUROSCI.3331-13.2014

Mukherjee, N., Wachutka, J., & Katz, D. B. (2019). Impact of precisely-timed inhibition of gustatory cortex on taste behavior depends on single-trial ensemble dynamics. Elife, 8. doi:10.7554/eLife.45968

Nakada, D., Saunders, T. L., & Morrison, S. J. (2010). Lkb1 regulates cell cycle and energy metabolism in haematopoietic stem cells. Nature, 468(7324), 653–658. doi:10.1038/nature09571

Nicolelis, M. A., Dimitrov, D., Carmena, J. M., Crist, R., Lehew, G., Kralik, J. D., & Wise, S. P. (2003). Chronic, multisite, multielectrode recordings in macaque monkeys. Proc Natl Acad Sci U S A, 100(19), 11041–11046. doi:10.1073/pnas.1934665100

Niell, C. M., & Stryker, M. P. (2008). Highly selective receptive fields in mouse visual cortex. J Neurosci, 28(30), 7520–7536. doi:10.1523/JNEUROSCI.0623-08.2008

Nolan, L. J., McCaughey, S. A., Giza, B. K., Rhinehart-Doty, J. A., Smith, J. C., & Scott, T. R. (1997). Extinction of a Conditioned Taste Aversion in Rats: I. Behavioral Effects. Physiology & Behavior, 61(2), 319–323. doi:10.1016/s0031-9384(96)00411-8

O’Brien, L. D., Wills, K. L., Segsworth, B., Dashney, B., Rock, E. M., Limebeer, C. L., & Parker, L. A. (2013). Effect of chronic exposure to rimonabant and phytocannabinoids on anxiety-like behavior and saccharin palatability. Pharmacol Biochem Behav, 103(3), 597–602. doi:10.1016/j.pbb.2012.10.008

Olshausen, B. A., & Field, D. J. (2004). Sparse coding of sensory inputs. Curr Opin Neurobiol, 14(4), 481–487. doi:10.1016/j.conb.2004.07.007

Piette, C. E., Baez-Santiago, M. A., Reid, E. E., Katz, D. B., & Moran, A. (2012). Inactivation of basolateral amygdala specifically eliminates palatability-related information in cortical sensory responses. J Neurosci, 32(29), 9981–9991. doi:10.1523/JNEUROSCI.0669-12.2012

Poo, C., & Isaacson, J. S. (2009). Odor representations in olfactory cortex: “sparse” coding, global inhibition, and oscillations. Neuron, 62(6), 850–861. doi:10.1016/j.neuron.2009.05.022

Rabiner, L. R. (1989). A tutorial on hidden Markov models and selected applications in speech recognition. Proceedings of the IEEE, 77(2), 257–286. doi:10.1109/5.18626

Richardson, R., Williams, C., & Riccio, D. C. (1984). Stimulus generalization of conditioned taste aversion in rats. Behavioral and Neural Biology, 41(1), 41–53. doi:10.1016/s0163-1047(84)90706-4

Sadacca, B. F., Mukherjee, N., Vladusich, T., Li, J. X., Katz, D. B., & Miller, P. (2016). The Behavioral Relevance of Cortical Neural Ensemble Responses Emerges Suddenly. J Neurosci, 36(3), 655–669. doi:10.1523/JNEUROSCI.2265-15.2016

Sadacca, B. F., Rothwax, J. T., & Katz, D. B. (2012). Sodium concentration coding gives way to evaluative coding in cortex and amygdala. J Neurosci, 32(29), 9999–10011. doi:10.1523/JNEUROSCI.6059-11.2012

Sakata, S., & Harris, K. D. (2009). Laminar structure of spontaneous and sensory-evoked population activity in auditory cortex. Neuron, 64(3), 404–418. doi:10.1016/j.neuron.2009.09.020

Shaw, R. J. (2009). LKB1 and AMP-activated protein kinase control of mTOR signalling and growth. Acta Physiol (Oxf), 196(1), 65–80. doi:10.1111/j.1748-1716.2009.01972.x

Viterbi, A. (1967). Error bounds for convolutional codes and an asymptotically optimum decoding algorithm. IEEE Transactions on Information Theory, 13(2), 260–269. doi:10.1109/tit.1967.1054010

Yamamoto, T., Azuma, S., & Kawamura, Y. (1984). Functional relations between the cortical gustatory area and the amygdala: electrophysiological and behavioral studies in rats. Exp Brain Res, 56(1), 23–31. doi:10.1007/BF00237438

Yamamoto, T., Fujimoto, Y., Shimura, T., & Sakai, N. (1995). Conditioned taste aversion in rats with excitotoxic brain lesions. Neuroscience Research, 22(1), 31–49. doi:10.1016/0168-0102(95)00875-t

Yirmiya, R., Lieblich, I., & Liebeskind, J. C. (1988). Reduced saccharin preference in CXBK (opioid receptor-deficient) mice. Brain Research, 438(1-2), 339-342. doi:10.1016/0006-8993(88)91360-1

Yiu, A. P., Mercaldo, V., Yan, C., Richards, B., Rashid, A. J., Hsiang, H. L., … Josselyn, S. A. (2014). Neurons are recruited to a memory trace based on relative neuronal excitability immediately before training. Neuron, 83(3), 722–735. doi:10.1016/j.neuron.2014.07.017

Yoshida, T., & Katz, D. B. (2011). Control of prestimulus activity related to improved sensory coding within a discrimination task. J Neurosci, 31(11), 4101–4112. doi:10.1523/JNEUROSCI.4380-10.2011

